# SMRT Genome Assembly Corrects Reference Errors, Resolving the Genetic Basis of Virulence in *Mycobacterium tuberculosis*

**DOI:** 10.1101/064840

**Authors:** Afif Elghraoui, Samuel J Modlin, Faramarz Valafar

**Affiliations:** Laboratory for Pathogenesis of Clinical Drug Resistance and Persistence, Biological and Medical Informatics Research Center, San Diego State University, San Diego, CA

## Abstract

The genetic basis of virulence in *Mycobacterium tuberculosis* has been investigated through genome comparisons of its virulent (H37Rv) and attenuated (H37Ra) sister strains. Such analysis, however, relies heavily on the accuracy of the sequences. While the H37Rv reference genome has had several corrections to date, that of H37Ra is unmodified since its original publication. Here, we report the assembly and finishing of the H37Ra genome from single-molecule, real-time (SMRT) sequencing. Our assembly reveals that the number of H37Ra-specific variants is less than half of what the Sanger-based H37Ra reference sequence indicates, undermining and, in some cases, invalidating the conclusions of several studies. PE_PPE family genes, which are intractable to commonly-used sequencing platforms because of their repetitive and GC-rich nature, are overrepresented in the set of genes in which all reported H37Ra-specific variants are contradicted. We discuss how our results change the picture of virulence attenuation and the power of SMRT sequencing for producing high-quality reference genomes.

---

Tuberculosis is a serious and pervasive public health problem [1]. It is a disease caused by infection of bacteria from the *Mycobacterium tuberculosis* complex (MTBC). The reference strain, *Mycobacterium tuberculosis* H37Rv, has an attenuated counterpart known as H37Ra that is available for studies where facilities to handle virulent samples are lacking. H37Ra exhibits a distinct colony morphology, an absence of cord formation, decreased resistance to stress and hypoxia, and attenuated virulence in mammalian models [2–4]. The H37Ra genome was assembled by Zheng and colleagues in 2008 and compared to H37Rv for the purpose of identifying the genetic basis of virulence attenuation [5]. The resulting sequence has been used as the primary avirulent reference genome for *M. tuberculosis* since its publication in 2008.

As genome sequencing technology has significantly improved [6], we sought to assess the ability of single-molecule, real-time (SMRT) sequencing for finishing mycobacterial genomes. In addition to a high overall GC-content, these genomes have GC-rich repetitive sequences, a source of systematic error for many sequencing protocols. Even sample preparation methods commonly used for shotgun Sanger sequencing are prone to such bias [7]. Sequencing errors in the H37Rv reference have been sought out, with some corrected, others remaining to be discovered, and still others discovered and remaining to be corrected [8,9]. The Pacific Biosciences RS II platform has been shown to produce finished-grade assemblies of microbial genomes exceeding the quality of Sanger sequencing [10–12].

In this study, we sequenced and assembled the genome of *M. tuberculosis* H37Ra and compared it to the reference sequence. We further compared both sequences against the reference sequence for *M. tuberculosis* H37Rv and re-evaluated the conclusions of Zheng and colleagues with respect to the genetic basis of virulence attenuation.

## Results

### Genome Assembly and Methylation Motif Detection

With the data from one sequencing run (SMRTCell), the genome assembled with 103x average coverage into a single contig containing 4426109 base pairs after circularization and polishing. Performing the assembly with data from two SMRTCells (217x average coverage) resulted in an identical sequence.

In the raw assembly, circularization was impeded by discrepancies in the edges of the contig, where an IS6110 insertion was present in only one of the two edges. It appears heterogeneously in our sample, as aligning our reads against our assembly shows that a minority of reads have interrupted mapping to this segment while the majority do not. With regard to base modifications, N6-methyladenine was detected in 99.67% of the instances of the partner sequence motifs CTGGAG and CTCCAG. The methylation of these motifs in both H37Ra and H37Rv was previously reported by Zhu and colleagues in H37Ra as part of their study of mycobacterial methylomes [13].

### Direct Comparison with the Hitherto H37Ra Reference Genome

Comparison of our assembly with the H37Ra reference sequence (NC 009525.1, hereafter referred to as H37RaJH, for Johns Hopkins) showed significant variation. We found 33 single nucleotide polymorphisms (SNPs), and 77 insertions and deletions in our assembly with respect to H37RaJH (Supplementary Data 1).

#### Structural Variations

Two of the insertions with respect to H37RaJH were substantial structural variations: one was an insertion of IS6110 into the gene corresponding to Rv1764 and the other was an in-frame insertion of 3456bp into the PPE54 gene.

The insertion of IS6110 into Rv1764 (an IS6110 transposase) is unsurprising, as IS6110 insert frequently into that general region of the genome, as well as within their transposase [14, 15]. This insertion was the heterogeneous insertion responsible for the discrepant contig ends in our raw genome assembly. Such heterogeneity implies either a lack of selection pressure on the insertion in culture, a recent emergence of the insertion, or both.

The 3456bp insertion in *ppe54* with respect to H37RaJH incidentally corresponds to a tandem duplication of a 1728bp sequence at the same site in H37Rv with 100% identity. The complete absence of this tandem repeat at this site in H37RaJH, however, is not necessarily an assembly error, as this is also observed in several clinical isolates (unpublished data). This, along with the 100% identity between each 1728bp duplicate of the tandem repeat with respect to H37Rv, lead us to believe that both the duplication in our sequence and the deletion observed in H37RaJH are instances of *in vitro* evolution, following the divergence of the lineages from which H37RaJH and our assembly were drawn.

These two structural variations, or, at least, very similar structural variations, have been observed previously in virulent strains of *M. tuberculosis*, and therefore likely do not contribute to virulence attenuation in H37Rv (unpublished data) [14,16], but shed light on *in vitro* evolution of this strain [8, 17].

### Analysis of Motif Variants in H37Ra and H37Rv

With the knowledge that the CTGGAG/CTCCAG motifs are methylated in both H37Ra and H37Rv [13], we determined the motif variants, or sequence polymorphisms that create or destroy motifs, between H37Rv and H37Ra. By first comparing H37RaJH to H37Rv, we see that all but two motif variants were due to structural variations. Both of these variants instantiate the CTGGAG motif in H37Ra where it is absent in the H37Rv reference sequence. The first is due to the *G* → *T* polymorphism at H37Rv position 2043284 (upstream of PPE30) in H37RaJH, but this variant is contradicted by our H37Ra assembly. The second is due to the *T* → *G* polymorphism at H37Rv position 2718852 (upstream of *nadD*) and confirmed by our H37Ra assembly, yet also appears in CDC1551 and is a previously reported sequencing error in H37Rv [8] that has not been applied to the current reference. Based on these results, DNA methylation and motif variants do not play a role in the attenuation of virulence in H37Ra.

### Status of Previously Reported “H37Ra-specific” Polymorphisms

With our assembly, we aimed to replicate the study performed by Zheng and colleagues when they first assembled the H37Ra genome [5]. In their study, they compared their assembly with H37Rv, then filtered out variants also present in CDC1551 (NC 002755.2) to find mutations likely specific to H37Ra [5]. Zheng and colleagues identified a set of mutations in H37Ra unique with respect to H37Rv and CDC1551 as “H37Ra-specific”. These mutations fall within or adjacent to (which we term “affecting”) 56 genes in H37Rv, which we refer to as the high-confidence (HC) gene set. While comparing the variants, Zheng and colleagues also discovered sequencing errors in the H37Rv reference sequence [5], a number of which were corrected in NC 000962.3 [9], the version used in our study.

To see how well the HC genes are supported by our assembly of H37Ra, we determined variants with respect to H37Rv for our assembly and H37RaJH and performed set comparisons after excluding mutations shared with CDC1551 (Supplementary Data 1-2). We then categorized the HC genes as follows. We labeled a gene “unsupported” if all mutations affecting it were observed only in H37RaJH. We labeled a gene “supported” if all mutations affecting it were observed in both H37Ra assemblies. Otherwise, we labeled a gene “adjusted” if it had a different variant profile between H37RaJH and our assembly in a manner distinct from the two categories defined above. Figure 1 shows example classifications based on these criteria.

**Figure 1:**
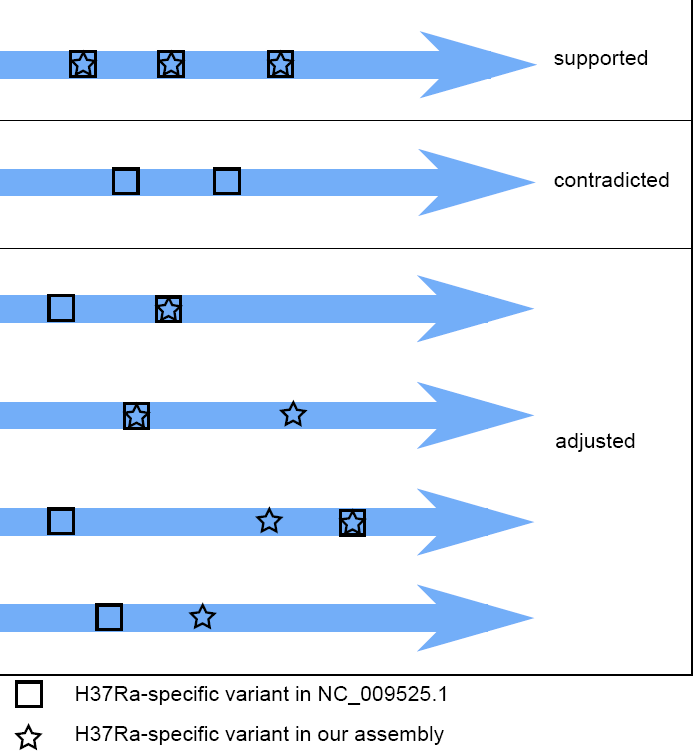
Example Classification of Genes Based on Variant Comparisons. Considering the profile of H37Ra-specific variants (those with respect to H37Rv not also appearing in CDC1551), a given gene (blue arrow) is categorized as “supported”, “contradicted”, or “adjusted” by our H37Ra assembly as a result of comparison with the hitherto reference sequence NC 009525.1. The illustration shows examples of the different variant profiles a gene could have and their resulting classifications. Genes in the “supported” and “contradicted” categories are strictly those where our assembly either fully matches the H37Ra reference (supported) or the H37Rv reference (contradicted). Multiple factors may cause a gene to be classified as “adjusted”. Such genes may have variant profiles not fully meeting the criteria of “supported” or “contradicted”, or they may have novel H37Ra-specific variants observed only in our assembly.

We first noted that two of the HC variants reported by Zheng and colleagues, those affecting *nadD* (Rv2421c) and *nrdH* (Rv3053), were included erroneously (Table 1d). These variants were a *T* → *G* mutation 44 bases upstream of *nadD*, at H37Rv position 2718852, and a 14bp deletion in the promoter of *nrdH*. These mutations, although confirmed by our assembly, also appear in CDC1551 and thus cannot be considered H37Ra-specific.

**Table 1:**
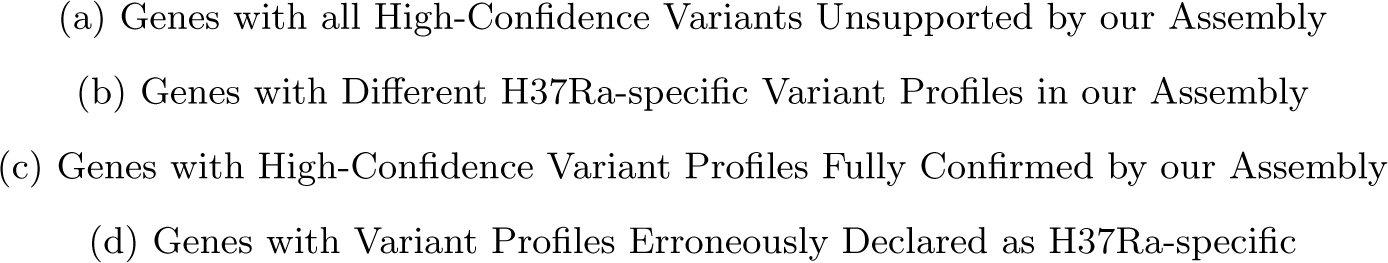
Status of Genes Previously Reported as Affected by H37Ra-specific Mutations.

Of the variants in the remaining 54 HC genes, our assembly contradicts 35 (Table 1a), adjusts 5 (Table 1b), and confirms 14 (Table 1c). We then considered how these results affect the picture of how the genotypic differences between H37Rv and H37Ra give rise to the phenotypic differences observed between the two strains, which are discussed below and depicted graphically in Figure 3. As our analysis focused on the HC gene set reported by Zheng and colleagues [5], we did not re-evaluate whether additional genes and variants should belong to this grouping. We did, however, carefully consider all variants unique to our assembly (Table 2) and their potential effect on the organism’s phenotype.

**Table 2:**
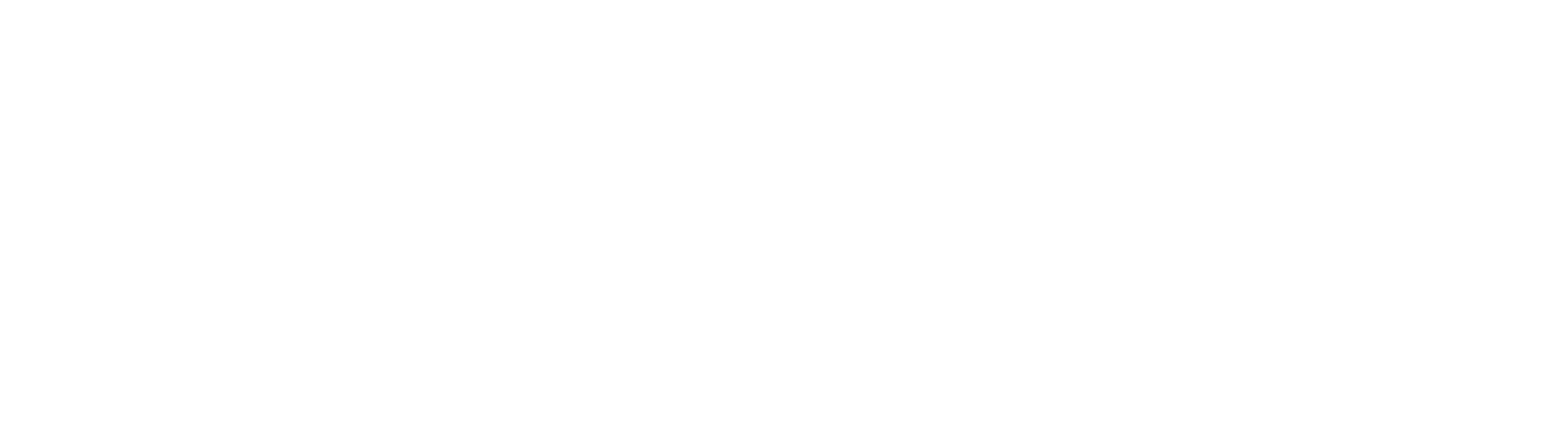
Variants in H37Ra Unique to our Assembly

#### Accuracy of the H37Rv Reference Sequence

Ioerger and colleagues listed 73 polymorphisms (excluding those in PE_PPE genes) with respect to the H37Rv reference shared between six H37Rv strains from different laboratories, but considered all but one of them as errors in the reference sequence because they also appeared in the H37Ra reference [8]. The remaining polymorphism was a *A* → *C* transversion at position 459399, a position upstream of Rv0383c masked by a 55bp deletion in H37RaJH. Interestingly, our assembly contradicts this 55bp deletion, but is in perfect concordance with the transversion at position 459399. The revelation that H37Ra is in fact the same as all H37Rv strains at this position invalidates the maximum parsimony tree in figure 1 of their publication [8]. Thus, through our improved assembly of the H37Ra genome, we have identified an additional error in H37Rv, the standard reference genome of *M. tuberculosis*.

#### SNPs Previously Reported to Cause Expression Changes in H37Ra are Contradicted by Our Assembly

Interestingly, SNPs in the putative promoter regions of two genes, *phoH2* and *sigC*, found by Zheng and colleagues to be up-regulated *in vitro* and down-regulated in macrophage in H37Ra relative to H37Rv, were contradicted by our assembly [5]. Zheng and colleagues attributed this differential expression to these (now contradicted) SNPs, but it appears there instead must be a distal causative factor driving the observed expression changes of both genes. The SNP affecting *sigC* has been cited as the cause of the differential expression of SigC in macrophages relative to H37Rv [18, 19], illustrating how incorrect SNPs Previously Reported to Cause Expression Changes in H37Ra are Contradicted by Our Assembly

#### SNPs Previously Thought to Affect Polyketide Synthesis in H37Ra are Contradicted by Our Assembly

Altered polyketide synthesis has been proposed as one of the primary mechanisms attenuating virulence in H37Ra, through disrupting phthiocerol dimy-cocerosate (PDIM) production, which has shown to manifest deleteriously in H37Ra [20,21]. Our assembly contradicts both reported SNPs in *pks12* (polyke-tide synthase 12) of H37RaJH. This means that some factor other than disruption of *pks12* causes the observed lowered PDIM production in H37Ra. Thus, it remains unclear which (epi)genomic factor(s) underlie the observed reduction in PDIM synthesis in H37Ra, as supported variants (those in *phoP* and *nrp*) once considered to cause this reduction [22] have been shown not to [23, 24]. However, it is possible the decreased production of PDIMs is merely an artifact of repeated subculturing *in vitro* [17].

#### Variants in *phoP*, *mazG*, and *hadC* Account for Much of the Virulence Attenuation in H37Ra

Of all the HC genes, only variants in *phoP*, *mazG*, and *hadC* have been connected strongly with virulence attenuation in H37Ra through wet-lab work, each of which our assembly supports.

Of these, the most thoroughly studied is the nsSNP (S219L) in the DNA-binding region of *phoP*, part of the two component *PhoPR* regulatory system. There is an abundance of literature linking *phoP* to virulence attenuation in H37Ra, through several mechanisms, including disrupted sulfolipid and trehalose synthesis (Figure 2), diminished ESAT-6 secretion, and additional downstream effects from altered expression of other genes under its regulon [5,18,23,25–30]. However, several of these studies also show that *phoP* alone [23,29] is not responsible for virulence attenuation in H37Ra, but rather that the genomic cause behind virulence attenuation in H37Ra is multifactorial.

**Figure 2:**
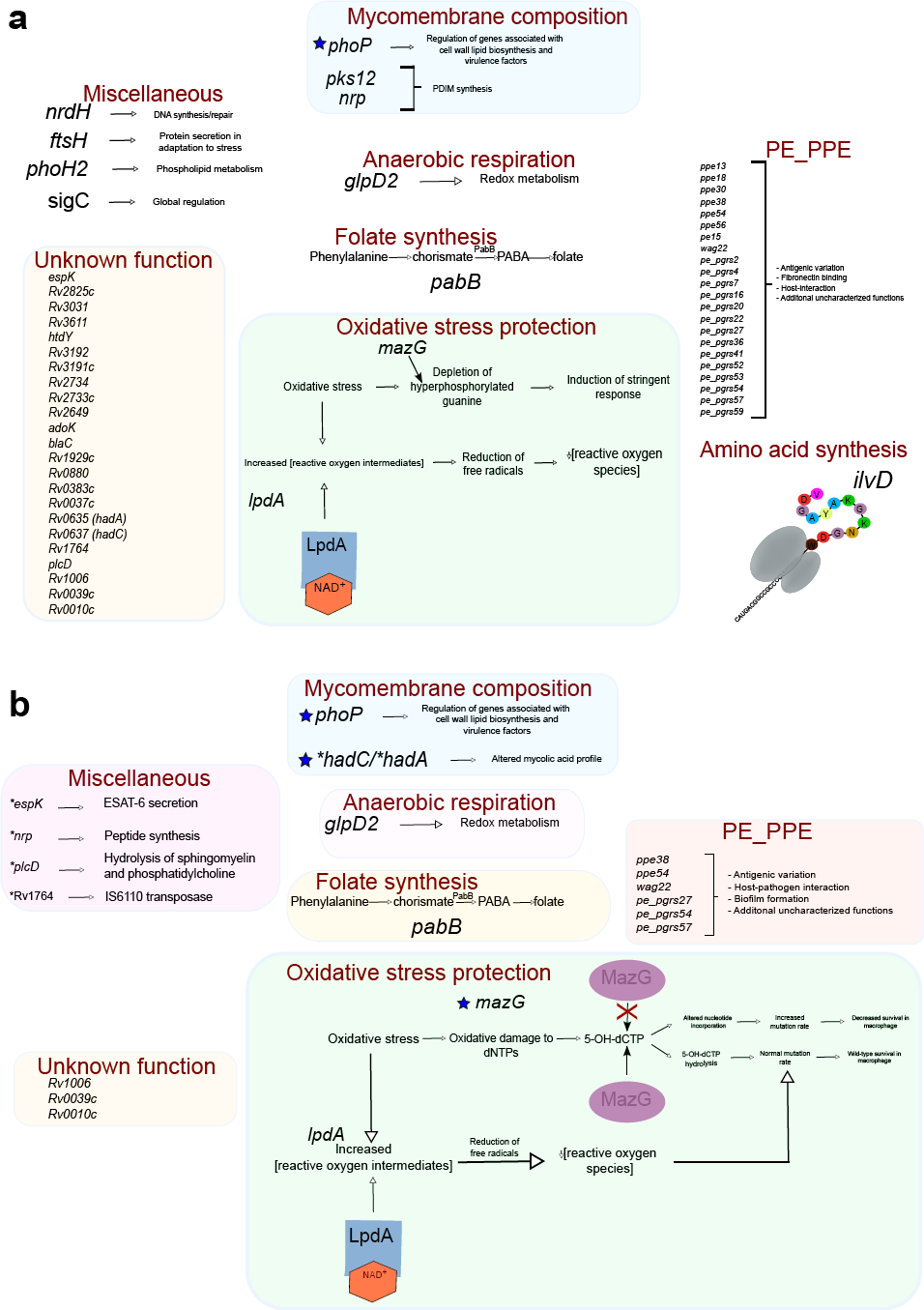
Cell Wall Differences in H37Ra and H37Rv. A) State of knowledge following publication of H37RaJH. At this time it was known that the SNP in the DNA-binding site of phoP abrogated synthesis of sulfolipids (yellow) and acyltrehaloses (purple and red) of the mycomembrane outer leaflet, while two SNPs in pks12, both of which were refuted in our assembly, were thought to cause the observed lack of phthiocerol dimycocerosates (blue) in H37Ra. B) Current state of knowledge. Advances were made in understanding the inner leaflet. A single nucleotide, frameshift deletion in the now annotated hadC gene was shown by Slama and colleagues [33] to alter the mycolic acid profile in three distinct ways: i. Lower proportion of oxygenated mycolic acids (K-MA and Me-MA; green and blue carbon skeletons, respectively) to α-MAs (orange carbon skeleton). There are seven Me-MAs depicted in H37Rv compared to three in H37Ra, reflecting the proportions reported by Slama and colleagues [33]. ii. Extra degree of unsaturation (red circles) in H37Ra mycolic acids due to truncation of the HadC protein in H37Ra. iii. Shorter chain lengths of mycolic acids in H37Ra. Note that Me-MAs have larger loops in H37Rv than in H37Ra, and that the height of the α-MAs is shorter in H37Ra than H37Rv. Carbon chain lengths are based on results reported by Slama and colleagues. The folding geometry of the mycolic acids is depicted in panel B, as described by Groenewald and colleagues [50], and inspired by the illustration style of Minnikin and colleagues [51].

**Figure 3:**
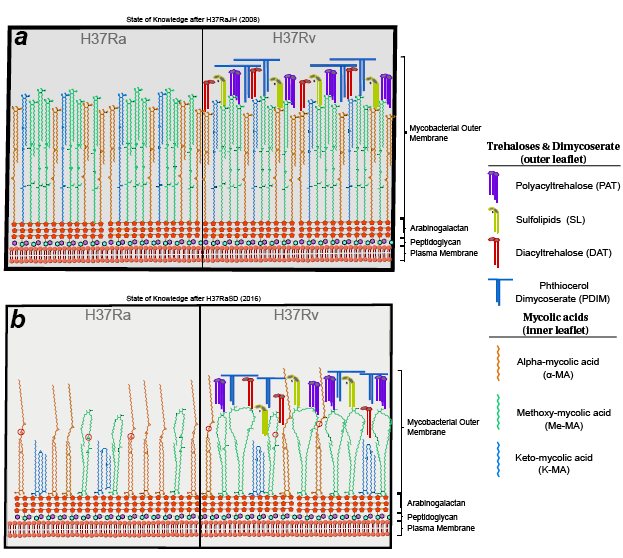
Visualization of the Reduced Set of H37Ra-specific Variants and Their Effect on Phenotype. Our assembly contradicts many variants previously thought to be H37Ra-specific, reducing the number of genes that may contribute to H37Ra’s virulence attenuation. Several of these genes have been reassigned function since the first published assembly of the H37Ra genome [5], which is reflected in the figure. Blue stars indicate that the H37Ra-specific variant(s) in that gene has been shown to confer a phenotypic change in H37Ra relative to H37Rv in wet-lab studies. For these genes, the mechanisms affected by the H37Ra-specific variant are illustrated in detail (see Figure 2 for hadC and phoP). For website genes, their general function is described or briefly illustrated. a) The set of genes identified to carry H37Ra-specific polymorphisms in the original H37Ra genome publication [5] and their contribution to phenotype as understood at that time. 57 genes are affected, the majority of which were PE_PPE genes or were of unknown function. b) The set of genes with H37Ra-specific variants confirmed by our assembly is reduced markedly, particularly in PE_PPE genes, highlighting the strength of single-molecule sequencing in resolving GC-rich and repetitive stretches of DNA. Genes with functions not yet characterized were also reduced significantly. Though in a few instances this was because these genes’ function was characterized between 2008 and now (indicated by an asterisk), most were due to our assembly showing that they matched that of H37Rv and, therefore, are not H37Ra-specific.

The second gene, *mazG*, has a nsSNP (A219E) in a region coding for a highly conserved alpha-helix residue in its protein product, a nucleoside triphosphate (NTP) pyrophosphohydrolase [5]. MazG exhibits diminished hydrolysis activity in H37Ra relative to both MazG in H37Rv and MazG of the fast-growing *M. smegmatis*. Wild-type MazG hydrolyzes all NTPs, including those that are mu-tagenic and appear more frequently with oxidative stress (Figure 3b), which is experienced by the bacterium inside activated macrophages [31]. This decreased ability to hydrolyze mutagenic NTPs contributes to virulence attenuation in H37Ra [32].

In the third gene, *hadC*, there is a frameshift-inducing 1-bp insertion, which creates a premature stop codon and results in truncation of HadC. *hadC* is a member of the essential *hadA-hadB-hadC* gene cluster, which forms two hy-dratases (HadAB and HadBC) of the *M. tuberculosis* fatty acid synthase II system. Our assembly and H37RaJH both show a 5-bp insertion in *hadA* which, along with *hadC*, are the only genes with variants in H37Ra [33] that encode proteins known to be necessary for mycolic acid synthesis.

Recent complementation and knockout studies using *hadC* from H37Ra and H37Rv showed that intact HadC is necessary for cord formation, and that the truncated form *H37Ra/hadC* affects length and oxygenation of mycolic acids (Figure 2b). Furthermore, when tested in murine lung and spleen, *H37Ra/hadC_Rv_* grew an intermediate amount of colony forming units, between that of H37Ra and H37Rv, at a level commensurate with *H37RvΔhadC* which suggests that the H37Ra *hadC* variant underlies some of its virulence attenuation [33].

Interestingly, while both our assembly and H37RaJH harbor a 5-bp insertion in *hadA*, sequences obtained by Lee, Slama, and their respective colleagues do not [29,33]. These two sequences were both derived from a culture from Institut Pasteur, while ours and that of Zheng and colleagues [5] were acquired directly from ATCC, which suggests that the two cultures diverged *in vitro* prior to sequencing despite sharing the same ATCC identifier. We expect the deleterious effects of *hadC_Ra_* shown by Slama and colleagues would be exacerbated by the 5bp insertion in our assembly, as it results in an abnormal HadAB enzyme which, when normal, has been posited to compensate for faulty HadBC [33]. However, the experiments discussed above indicate that the *hadC* variant alone is sufficient to attenuate virulence, and is one of the primary sources of attenuation in H37Ra.

#### Copy Number Variation in lpdA Promoter

The polymorphism reported in H37RaJH that affects *lpdA* (NAD(P)H quinone reductase) is a third (as opposed to the two in H37Rv) 58bp repeat in its promoter region. Our assembly reveals an additional two copies of this 58bp region, resulting in a total of five copies of the repeat. LpdA has been shown to protect bacilli from oxidative stress and improve *M. tuberculosis* survival in a mouse model, which suggests that if this copy number variation disrupts typical expression of LpdA, it may contribute to the phenotype of H37Ra [34]. This may also affect the expression of *glpD2* (glycerol-3-phosphate dehydrogenase), as it shares an operon with *lpdA* [5].

#### Variants Affecting Uncharacterized Hypothetical Genes

Several genes classified with unknown or hypothetical functions were among the HC genes of H37RaJH (Table 1). Our assembly contradicts all variants in the majority of these, leaving three which we supported in full.

Though none of these genes have an implicated role in virulence in the literature, they may in reality. These genes should be investigated, as they are three of the few supported HC genes yet to be explored. The value of exploring hypothetical genes is evidenced by the recent discovery of a significant contribution of HadC [33]—the function of which was unknown when H37RaJH was published—to virulence attenuation in H37Ra (Figure 3).

#### Significant Reduction of H37Ra-specific Variants in PE_PPE genes

The PE_PPE family of genes is unique to mycobacteria but poorly characterized, both functionally and genomically, in *M. tuberculosis*, the latter owing to the family’s high-GC content and repetitive nature [35]. Evidence for contribution from PE_PPE family members to virulence has amassed support since 2008 [36–39], but this gene family was the most drastically altered by our assembly: while PE_PPE genes comprise approximately 10% of the genome, they account for nearly half (16/35) of the unsupported genes. It is likely that the majority of these are errors in H37RaJH rather than manifestations of hypervariability, as few PE_PPE genes fell into the adjusted or novel categories, as one would expect if they were due to hypervariability.

Consequently, some extant work examining polymorphic PE_PPE genes between H37RaJH and H37Rv is invalidated by our assembly. For example, our assembly contradicts or changes the variant profile of all four PE_PPE genes reported to be positively selected for in H37Ra in an evolutionary genomics study by Zhang and colleagues [38] using H37RaJH.

Another study affected profoundly by our results is that of Kohli and colleagues [36], which used H37RaJH and H37Rv in an *in silico* genomic and proteomic comparison of PE_PPE family genes. Though our assembly renders much of the results from their analyses invalid, applying their methodology to our updated assembly would yield interesting results.

Our assembly contains polymorphisms in 6 of 22 genes that encode PE_PPE family members reported as unique to H37RaJH (Table 1, Figure 3b). Of the three PE_PPE family members fully corroborated by our sequence, one was the duplication of *ppe38*, which McEvoy and colleagues have also identified in 3 different samples of H37Rv, suggesting this duplication likely plays no role in virulence [40]. All 3 of the adjusted PE_PPE family members, as well as the supported *Wag22* and *PPE13*, belong to PE_PPE sublineage V. Sublineage V members comprise the majority of PE_PPE proteins that interact with the host, and are overrepresented in proteomic studies of *in vivo* infection [35]. This enrichment of subfamily V PE_PPE family members in the set of supported or adjusted genes suggests they may be more integral to virulence attenuation in H37Ra than other PE_PPE family members. The role of PE_PPE family members in virulence should become better understood as more genomes are sequenced using third-generation platforms.

In addition to the differences due to sequence alterations in PE_PPE family genes, the corroborated polymorphism in *phoP* may confer altered expression of many PE_PPE family proteins, as at least 13 are under its regulon [35], which could mediate some virulence attenuation.

The precise roles of PE_PPE family members have yet to be elucidated in full. It is difficult to evaluate rigorously the effect of each PE_PPE variant, as their function in wild-type *M. tuberculosis* is poorly characterized [35]. Moreover, their contribution to virulence may well require complexities of the native host environment beyond what can be replicated *in vitro* or *ex vivo* with current technology. Thus, the role the polymorphisms in this family play in the phenotype of H37Ra compel further research, which our reduction of variants has made more tractable.

## Discussion

Since its publication in 2008 [5], several studies have used the whole genome [8, 36, 41–46] of H37RaJH, or the reported differences between H37RaJH and H37Rv [47] in their analyses. Our improved assembly changes the implications of several of these *in silico* analyses. Additionally, several studies have used the set of genes with variants in H37RaJH with respect to H37Rv to guide wet-lab experiments [48, 49]. Re-examining these studies with our assembly of H37Ra may yield novel insights, as unsupported variants can serve as a retroactive control.

Our *de novo* assembly using single-molecule sequencing has reduced the set of genes polymorphic to H37Rv by more than half, clarifying which genomic factors most likely give rise to virulence attenuation and other H37Ra-specific phenotypes. For an expanded discussion of genes affected and their ties to virulence, see the supplementary note. Supported variants affecting PhoP, MazG, and HadC have been experimentally affirmed [23, 32, 33], gaining insight into how they manifest in the phenotype of H37Ra, but basic mechanisms for their contributions are not fully elucidated. A few other supported or adjusted genes (*lpdA, pabB, and nrp*) have been indirectly connected to the avirulence of H37Ra through experiments on other mycobacterial species [22] or H37Ra complementation studies measuring proxies of virulence [48].

It is clear that the nsSNP in *phoP* remains a potent mediator of virulence of *M. tuberculosis* through affecting SL and ATHL activity (Figure 3), while the truncation of HadC enfeebles the mycomembrane (Figure 3b). Polymorphisms in *mazG* and *lpdA* may each confer compromised stress response mechanisms in H37Ra (Figure 3), which are critical to enduring the intramacrophage environment of the host [32, 34]. Variants affecting genes with regulatory functions— *phoP* and others with roles in regulation not yet known—may also cause downstream effects on H37Ra phenotype, which may prove difficult to characterize. The variants in genes of the PE_PPE family and hypothetical genes (Rv0010c, Rv0039c, and Rv1006) potentially contribute to virulence attenuation through mechanisms not yet identified. Thus, with the greater accuracy of our assembly, wet-lab studies can focus on the true differences between the H37Ra and H37Rv, and computational studies will be in greater concordance with reality, yielding more useful results.

The advantages of single-molecule sequencing are readily apparent in our results. The random error profile of this technology allows for consensus accuracy to increase as a function of sequencing depth [10]. Performing the assembly with a doubled sequencing depth resulted in an identical sequence, indicating that we were able to maximize the sequence’s accuracy with a single sequencing run. The long reads produced by this technology allowed us to easily and unambiguously capture known structural variants in H37Ra, as well as two novel to the strain. We were also able to fully resolve the GC-rich and repetitive PE_PPE genes, sequences which compound the weaknesses of most sequencing technologies. As a result, our assembly demonstrates that H37Ra is significantly more similar to H37Rv than indicated by H37Ra’s Sanger-based reference sequence, with contradicted variants overrepresented in the difficult sequences of the PE_PPE genes. While *in vitro* evolution may underlie some of the differences between our assembly and H37RaJH, we believe that most of the contradicted variants (Table 1a) reflect sequencing errors in H37RaJH due to the disparity in sequencing quality. Regardless, the contradicted variants should not be considered as characteristic of H37Ra or its attenuated virulence. These sites were concordant with H37Rv and we did not find additional polymorphic PE_PPE genes with respect to H37Rv (Table 2), indicating a disparity in sequencing accuracy even among the Sanger-based references. On the other hand, the fact that we have not resequenced H37Rv and CDC1551 is a limitation of our study, where we have relied on their Sanger-based reference sequences for determining H37Ra-specific variants. We believe that the impact of the latter is minimal and that the former is the dominant factor, considering the level of concordance with H37Rv and CDC1551 in the instances where our sequence disagreed with H37RaJH.

Studies that have relied on the accuracy of PE_PPE sequences in the H37Ra reference sequence were the most severely impacted by our study. We consequently advise caution when analyzing GC-rich and repetitive sequences among reference genomes, not to mention draft genomes. As *de novo* assembly can be routinely performed for microbes using single-molecule sequencing, we strongly recommend this for mycobacteria, especially because of their PE_PPE genes.

## Competing interesst

The authors declare that they have no competing interests.

## Author’s contributions

F.V., A.E., and S.J.M. designed the study. A.E. performed the *de novo* assembly, methylation analysis, and comparative genomics analyses. S.J.M. performed the literature review, interpreted the results, and wrote the supplementary note. A.E. and S.J.M. prepared the manuscript, which was reviewed and approved by all authors.

## Acknowledgements

We would like to thank Jason Chin and Richard Hall from Pacific Biosciences for discussions on *de novo* assembly methodology and quality assessment. Logan Fink also provided some assistance with quality assessment of the assembly. We would also like to thank Antonino Catanzaro, Timothy Rodwell, and their staff for bacterial culture and DNA extraction. Jonas Korlach, Anthony Baughn, Sarah Ramirez-Busby, Ragavi Shanmugam, Carmela Chan, Amy Goodmanson, Daeheon Oh, and Logan Fink reviewed and provided helpful feedback on drafts of the manuscript. This work was funded by a grant from National Institute of Allergy and Infectious Diseases (NIAID Grant No. R01AI105185). A.E., S.J.M., and F.V. were supported by this grant. S.J.M. was also supported by scholarships from a National Science Foundation Grant (no. 0966391).

## Supplementary Information

### Supplementary Note

**Expanded Discussion of Virulence Attenuation Mechanisms in *M. tuberculosis* H37Ra**

### Supplementary Data 1

**Raw Variants**

Zip archive containing the following data in Variant Call Format (VCF):

**A6_7-H37Ra_NC009525_1.vcf** Variants in our H37Ra assembly with respect to the H37Ra reference sequence.

**A6_7-H37Rv_NC000962_3.vcf** Variants in our H37Ra assembly with respect to the H37Rv reference sequence.

**H37Ra_NC009525_1-H37Rv_NC000962_3.vcf** Variants in the H37Ra reference sequence with respect to the H37Rv reference sequence.

### Supplementary Data 2

**Annotated Variants with Respect to H37Rv**

Spreadsheet containing annotated variants in our assembly and the H37Ra reference sequence with respect to the H37Rv reference sequence. The sheets separate variants that are common to the two H37Ra assemblies and those that are unique to each.

### Supplementary Data 3

**Computer Code used for Analyses**

## Online Methods

### Sample Preparation and Whole-Genome Sequencing

*M. tuberculosis* H37Ra (ATCC25177) was obtained from ATCC and cultured on Lowenstein-Jenson slants and Middlebrook 7H11 plates. Cultures were incubated until growth of a full bacterial lawn. DNA was extracted using Genomic-tips (Qiagen Inc.) following the manufacturer’s sample preparation and lysis protocol for bacteria with the following modifications. Culture was harvested directly into buffer B1/RNAse solution, homogenized by vigorous vortex mixing and inactivated at 80°C for 15 minutes. Lysozyme was added and incubated at 37°C for 30 minutes followed by the addition of proteinase K and further incubation at 37°C for an additional 60 minutes. Buffer B2 was added and the mixture was incubated overnight at 50°C. Wide-bore pipet tips were used to optimize recovery of large DNA fragments. The remainder of the Genomic-tip protocol was carried out exactly as described by the manufacturer. DNA purity and concentration was analyzed on a Nanodrop 1000 (Thermo Scientific). The DNA was then sequenced using two SMRTCells on the Pacific Biosciences RS II instrument with the P6-C4 chemistry and a 20kb insert library preparation.

### Genome Assembly and Methylome Determination

The genome was assembled using Pacific Biosciences’ Hierarchical Genome Assembly Process [12] (HGAP) as implemented in SMRTAnalysis 2.3.0. This version of SMRTAnalysis provides two implementations of HGAP: HGAP.2 and the newer HGAP.3. HGAP.3 differs from HGAP.2 by replacing the Celera Assembler’s assembly consensus step with Pacific Biosciences’ speed-optimized implementation. We used HGAP.2 because, in our experiments, we found that HGAP.3 consistently produced spurious contigs while HGAP.2 did not.

The overlapping ends of the contig, an artifact of the assembly due to the circularity of the chromosome, were trimmed and joined using the minimus2 program from AMOS (http://amos.sourceforge.net). Discrepancies between the contig ends were resolved manually by selecting an authoritative sequence and trimming the discrepant one. The circularization was also performed with Circlator [52] to confirm the minimus2 results.

We validated the assembly structure using PBHoney [53], a structural variation detection tool, by using our assembled genome as input. Any structural variations detected would indicate potential misassemblies.

Circularization was followed by three rounds of assembly polishing using Quiver in SMRTAnalysis. Quiver was used with the maximum coverage parameter set to 1000 and otherwise default settings.

The methylome was determined using the base modification and motif analysis protocol in SMRTAnalysis.

### Comparative Genomics

In all cases, variants were determined using GNU diff (http://www.gnu.org/software/diffutils), an implementation of Myers’ algorithm for solving the longest-common-subsequence problem [54, 55] and converted to the Variant Call Format for analysis. This process is implemented in our custom tool, biodiff (http://www.github.com/valafarlab/biodiff). Because insertions and deletions in repetitive regions can be represented equivalently in multiple ways, we normalized the variants using the “norm” function of bcftools (http://samtools.github.io/bcftools), giving every mutation a standard representation to facilitate a proper comparison. Variants were then compared using bcftools isec and annotated using the Ensembl Variant Effect Predictor [56]. Motif variants were analyzed using *in villa* code.

### Literature Review

In order to gain a holistic view of the research built off of and conclusions drawn from the unique variants of H37RaJH with respect to H37Rv, we performed an exhaustive literature review. Common names and Rv numbers were searched using Google scholar within all publications which cited Zheng et al, 2008 [5] as of March 14th, 2016, for all genes with H37RaJH specific variants within CDS or potential promoter regions, according to Table 2 of [5]. All mentions of these genes were compiled and evaluated to illustrate how our assembly alters the picture of how the genomic differences between the reference strains contribute to the observed virulence attenuation of H37Ra (Discussion). Genes are discussed in the present study according to the H37Rv annotation (as opposed to H37Ra’s own annotation), as this convention relates to extant publications most readily.

### Data Availability

The assembled sequence and raw sequencing data for this project are available through NCBI under Bioproject PRJNA329548.

### Code Availability

Our motif variants detection tool is available from http://github.com/valafarlab/motif-variants. Analysis code for this study is provided as Supplemental Data 3.

